# Unique Pakistani gut microbiota highlights population-specific microbiota signatures of type 2 diabetes mellitus

**DOI:** 10.1101/2022.06.23.497392

**Authors:** Afshan Saleem, Aamer Ikram, Evgenia Dikareva, Emilia Lahtinen, Dollwin Matharu, Anne-Maria Pajari, Willem M. de Vos, Fariha Hasan, Anne Salonen, Ching Jian

## Abstract

Biogeographic variations in the gut microbiota reflect host and environmental factors delineating human populations, and are pivotal to understand global patterns of host-microbiota interactions in health and prevalent lifestyle-related diseases, such as type 2 diabetes mellitus (T2D). Pakistani adults, having an exceptionally high prevalence of T2D, are one of the most understudied populations in microbiota research to date. The aim of the present study is to examine the gut microbiota across individuals from Pakistan and other populations of non-industrialized and industrialized lifestyles with a focus on T2D. The fecal samples from 94 urban-dwelling Pakistani adults with and without T2D were profiled by 16S ribosomal RNA gene amplicon sequencing and qPCR, and plasma samples quantified for circulating levels of lipopolysaccharide binding protein (LBP) and the activation ability of Toll-like receptor (TLR)-signaling. Publicly available datasets generated with comparable molecular methods were retrieved for comparative analysis. Overall, urbanized Pakistanis’ gut microbiota was similar to that of transitional or non-industrialized populations, depleted in *Akkermansiaceae* and enriched in *Prevotellaceae* (dominated by the non-Westernized clades of *Prevotella copri*). The relatively high proportion of *Atopobiaceae* appeared to be a unique characteristic of the Pakistani gut microbiota. The Pakistanis with T2D had elevated levels of LBP and TLR-signaling in circulation as well as gut microbial signatures atypical of other populations e.g., increased relative abundance of *Libanicoccus/Parolsenella*, limiting the inter-population extrapolation of gut microbiota-based classifiers for T2D. Taken together, our findings call for more global representation of understudied populations to extend the applicability of microbiota-based diagnostics and therapeutics.

## Introduction

The human gut microbiota (i.e., collection of microbes living in the gut) has been associated with various intestinal and extraintestinal diseases due to its considerable contribution to immune and metabolic homeostasis.^1^ Substantial biogeographic variations have been documented in the human gut microbiota, reflecting differences in lifestyle practices, hygiene and pathogen load, diet, medication as well as host genetics that altogether influence the assembly and composition of the gut microbiota.^2^ Studies have begun to unravel the impact of these biogeographic variations on health, potentially linking the evolution of the gut microbiota to the varying burden of non-communicable diseases and the prevalence of infectious diseases in different populations.^2^ Currently, more than two thirds of public human microbiota datasets originate from Europe and North America, whereas a large number of uncultured bacterial species that may have ramifications in health and disease exists in the understudied populations.^4, 5^ On the other hand, population studies indicate that the gut microbiota varies between individuals of similar^6, 7^ or different ethnicities^8–10^ residing in geographical proximity. Altogether, these intra- and inter-population differences limit the applications of microbiota-based diagnostics and treatments,^7^ which can be exemplified by the debate on microbiota signatures of type 2 diabetes mellitus (T2D). The pathophysiology of T2D, characterized by dysregulated glucose metabolism and insulin resistance, has been suggested to have a microbiota component via low-grade inflammation initiated by pathogen-associated molecular patterns (PAMPs; e.g., lipopolysaccharide (LPS)) and altered levels of short-chain fatty acids (SCFAs; primary saccharolytically-derived microbial fermentation products);^1^ both mechanisms were predominately inferred from preclinical animal models and associative studies from a few extensively-studied populations. However, recent studies have found little convergence of the microbiota characteristics of T2D across geographical regions^11^ or across cohorts of colocalized individuals of the same ethnicity.^12^

Pakistan as the world’s fifth-most populous country had the highest age-adjusted comparative diabetes prevalence in 2021;^13^ Pakistan is experiencing rapid urbanization with increasing consumption of the Western diet and lifestyle changes especially in the urban areas,^14^ which all have implications for the gut microbiota. However, virtually nothing is known about the gut microbiota of Pakistanis in a global context, as the population of 220 million is represented by < 100 samples from few pilot studies.^14, 15^ In general, Southern and central Asia is the most underrepresented region for microbiota research.^3^ To bridge the knowledge gap, we aimed to profile the gut microbiota of Pakistani adults with and without T2D, and to assess the generalizability of the gut bacterial signature of T2D across different cohorts. We systematically compared the healthy Pakistani microbiota to that of industrialized (represented by China, Japan and Finland) and transitional or non-industrialized (represented by Indonesia, Sudan and rural India) populations as well as that of Pakistani adults with T2D. Potential differences in the butyrate production capacity of the gut microbiota, circulating levels of lipopolysaccharide binding protein (LBP) and Toll-like receptor (TLR)-signaling in circulation between Pakistani participants with and without T2D were analyzed to gain mechanistic insights. We then evaluated whether a classification model of T2D based on the Pakistani gut microbiota can be extrapolated to other populations. Only publicly available datasets targeting the V3-V4 region of the 16S rRNA gene and employing the Illumina sequencing technology were included in the present study to minimize methodological artifacts, as previous studies suggest that using matching variable regions of the 16S rRNA gene represents one of the most essential factors for cross-study comparability.^16, 17^

## Results

### Taxonomic characteristics of the Pakistani gut microbiota in a global context

Our Pakistani cohort included 94 urban-dwelling adults living in and around the capital regions with and without confirmed T2D. Their gut microbiota was profiled using 16S rRNA gene amplicon sequencing targeting the V3-V4 region, which resulted in 17,160 ± 910 quality-controlled and chimera-checked reads per sample. Overall, Firmicutes and Actinobacteria were the most abundant phyla in the Pakistani population accounting for 63.7% and 25.2% of the total read counts on average, respectively, and they were observed in all the samples. Bacteroidetes and Proteobacteria constituted 5.3% and 4% of the total read counts, respectively, yet these microbes were found in 87% and 88% of the study participants, respectively. Verrucomicrobia (genus *Akkermansia*) made up less than 1% of the gut bacterial community and was detectable in only 16% of the individuals. Spirochaetes (dominated by *Spirochaetaceae/Treponema* 2), a phylum observed mainly in ancient and non-industrialized societies but absent in industrialized populations,^18^ was detected in 10 Pakistani participants with an average abundance of 0.13%.

Besides a pilot study showing a few phylum-level differences when comparing urban Pakistani adults’ gut microbiota to the now defunct uBiome database of unspecified origin,^14^ the urban Pakistanis’ gut microbiota has not been evaluated on a global scale. Therefore, we compared the healthy participants’ microbiota to their counterparts from publicly available datasets generated with comparable molecular methods for profiling the fecal microbiota, including an external cohort of Pakistanis sampled from the same area (Table S1). Visualization of the Bray-Curtis dissimilarity measure using Principal Coordinates Analysis (PCoA) revealed a separation of industrialized populations (China, Japan and Finland) from the rest along the primary axis (P = 0.001, PERMANOVA; Fig. 1A). Moreover, the East Asian populations clustered tightly, slightly separated from the Finnish cohort, mirroring a recent study showing the impact of East Asian ethnicity on the gut microbiota.^10^ The overall structure of the Pakistani gut microbiota in our cohort (Pakistan1/PAK1) overlapped with that of the Indonesian gut microbiota, but somewhat differed from the other Pakistani cohort (Pakistan2/PAK2). The rural Indian population occupied a separate corner in the ordination space. The two Pakistani cohorts showed no difference in microbiota α-diversity metrics, and generally were on the similar levels of that of Chinese and Japanese populations that had the lowest α-diversity (Fig. 1B). The Indian cohort had the highest microbiota α-diversity as previously reported for other non-industrialized populations.^2^ In terms of taxonomic composition of the gut microbiota on the family level, stark differences between the industrialized and transitional/non-industrialized populations existed in *Prevotellaceae* and *Akkermansiaceae; Prevotellaceae* was relatively dominant in both Pakistani cohorts and *Akkermansiaceae* represented as an absentee, similar to other non-industrialized populations (Fig. 1C). High prevalence of *Atopobiaceae* (dominated by *Olsenella* and *Libanicoccus*) was uniquely observed in the Pakistani gut microbiota. Other taxonomic features commonly found in both Pakistani cohorts included highly abundant *Bifidobacteriaceae* (dominated by *Bifidobacterium*) and *Erysipelotrichaceae* (dominated by *Catenibacterium* and *Holdemanella*) as well as exceptionally high proportions of *Enterobacteriaceae* and *Streptococcaceae* in some individuals (Fig. 1C). *Spirochaetaceae* (belonging to Spirochaetes) could be detected only in the Pakistanis (11% prevalence in PAK1 and 15% in PAK2), Sudanese (31% prevalence), and Indians (7% prevalence).

**Figure 1.**
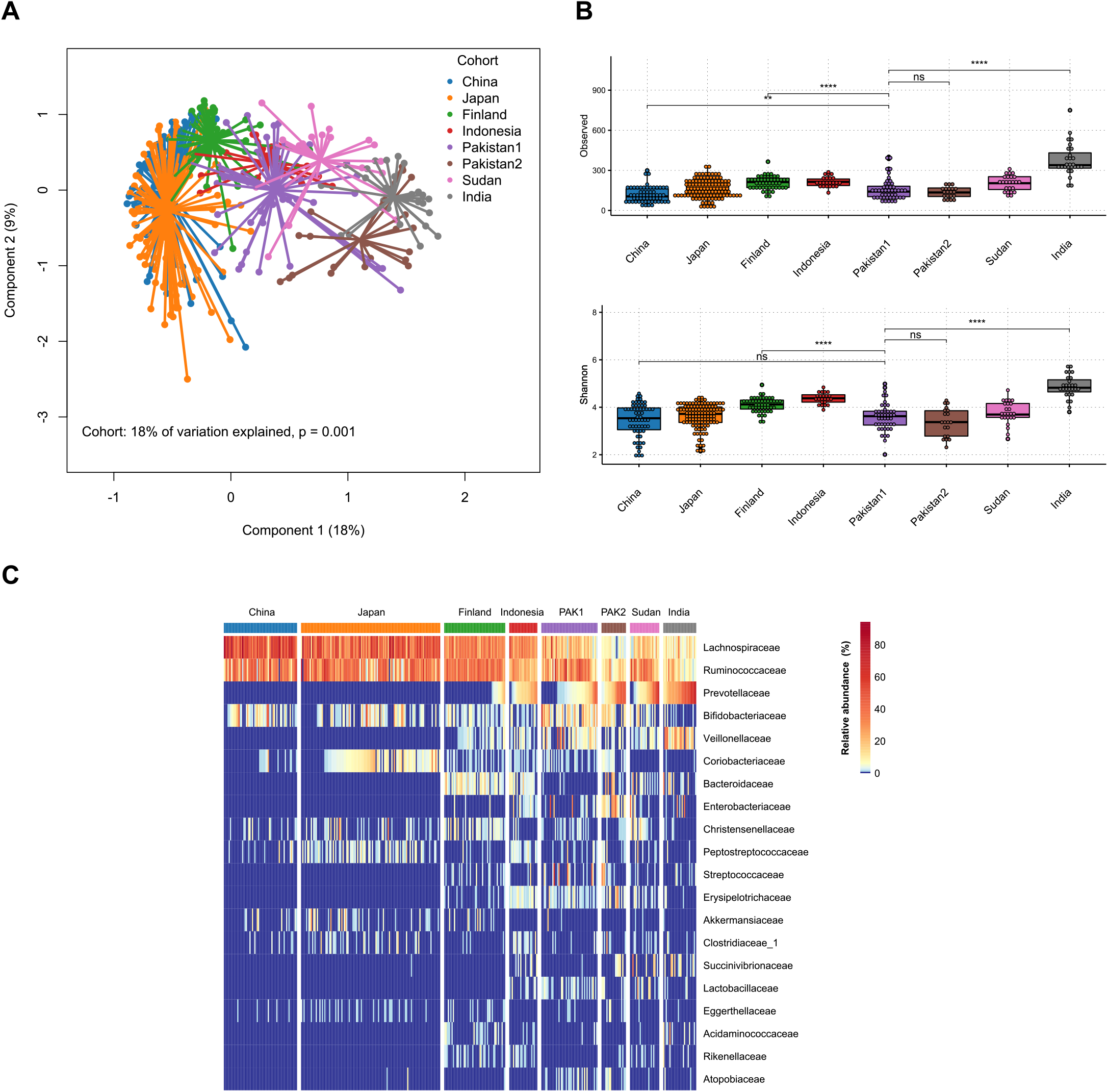
(**A**) Principal coordinate analysis (PCoA) plot of microbiota variation from different cohorts of healthy adults based on the Bray-Curtis dissimilarity matrix. Cohort explained 18% of the microbiota variation (P = 0.001, PERMANOVA). **(B)** Box plots showing microbiota α-diversity (observed richness and Shannon’s diversity) per each cohort. The center line denotes the median, the boxes cover the 25th and 75th percentiles, and the whiskers extend to the most extreme data point, which is no more than 1.5 times the length of the box away from the box. Points outside the whiskers represent outlier samples. Significance is calculated using the Wilcoxon rank-sum test. **** P < 0.0001; *** P < 0.001; ** P < 0.01; * P < 0.05; “ns” P > 0.05. (**C**) Heat map showing the relative abundance of the 20 most dominant bacterial families per cohort.

*Prevotella copri* of the *Prevotellaceae* family can be grouped into four clades (A, B, C, and D) depending on their genetic structure; clade A was ubiquitously found among both the Westernized and non-Westernized populations, while clades B, C, and D were predominantly found in non-Westernized populations.^19^ As the urban Pakistanis’ gut microbiota appears to exhibit features of the non-industrialized gut microbiota, we evaluated whether the urban Pakistani population has *P. copri* clades typical of non-Westernized populations. Consequently, we found that *P. copri* ASVs in both Pakistani cohorts belonged to clades B and C (Table S2).

### Taxonomic, functional and molecular signatures in the gut microbiota of Pakistanis with T2D

Focusing on our PAK1 cohort, age and sex were similarly distributed for the T2D patients and healthy controls, and for the patients prescribed with metformin and non-metformin hypoglycemic treatment. BMI, total cholesterol (TC), triglycerides (TG) and HbA1c were significantly higher in those with T2D or the non-metformin users (P < 0.05, Table 1). The individuals with and without T2D had significant differences in self-reported diets. Among the variables mentioned above, triglycerides and HbA1c significantly explained the variation in the gut microbiota in the model as determined by *EnvFit* (P < 0.01, Table 1).

**Table 1.**
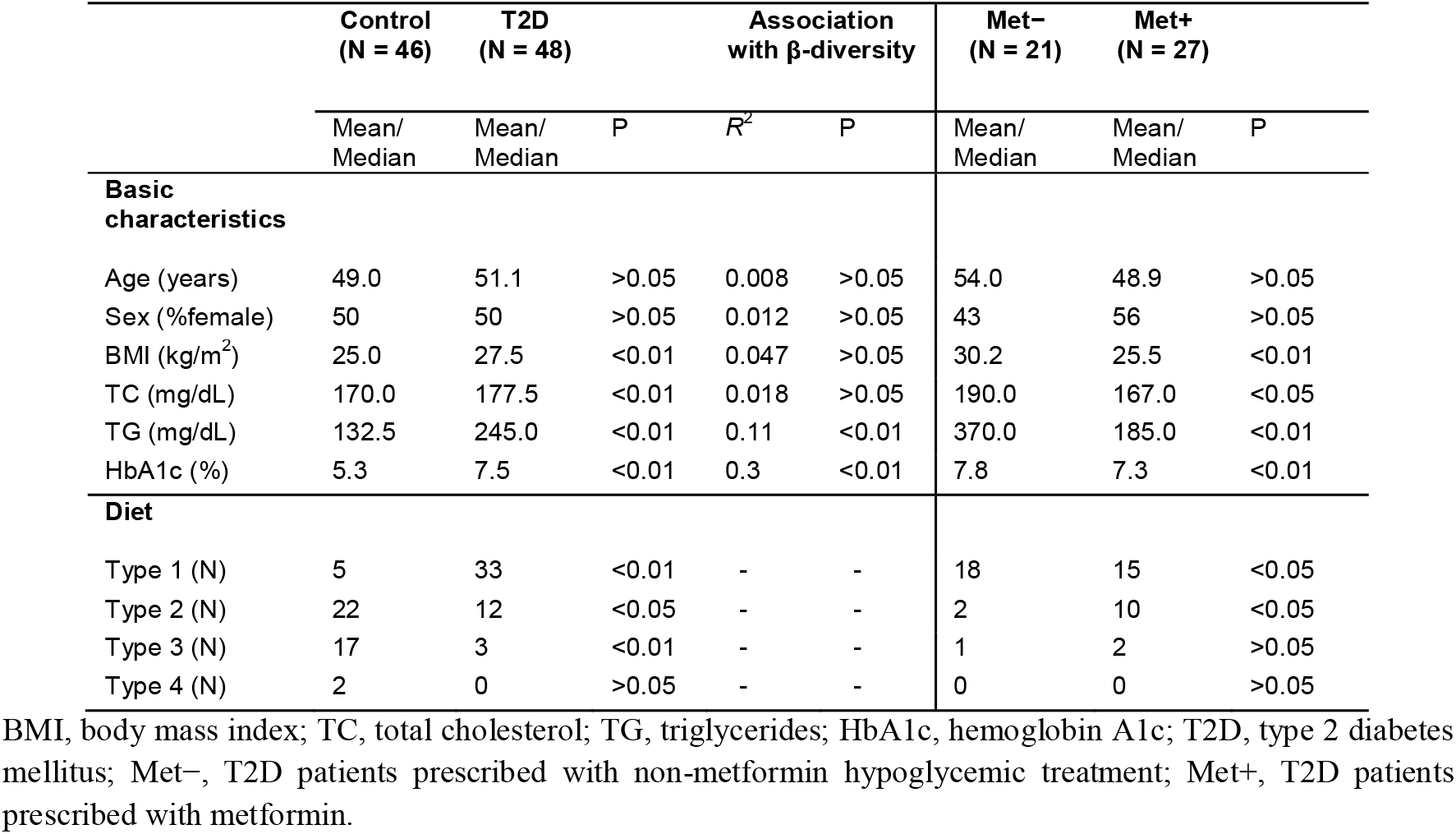
Anthropometric, biochemical parameters and dietary characteristics of participants, and their associations with the gut microbiota based on β-diversity.

Principal coordinate analysis (PCoA) showed a clear separation between the gut microbiota of patients and controls (Fig. 2A), demonstrating that T2D status was associated with the gut microbiota composition, explaining 9% of the total variation (P < 0.001, PERMANOVA). To identify taxa and functional modules driving the differences, we compared the abundance of individual bacterial families, genera, species and imputed KEGG pathways between the patients and controls using a negative binomial generalized linear model, as implemented in differential expression analysis for sequence count data version 2 (*DESeq2*). The analyses were additionally controlled for BMI and diet to disentangle their potential confounding effects. Striking differences (all FDR-P < 0.05) between the groups were found on all taxonomic ranks as well as in the functional modules with and without controlling for covariates. On the family level (Fig. 2B), *Lactobacillaceae* and *Coriobacteriaceae* were enriched and *Ruminococcaceae* depleted in T2D patients, which remained significant after adjusting for BMI and diet. Moreover, a marked decrease in *Prevotellaceae* (mainly attributable to *Prevotella 9/Prevotella copri*) was noted, which lost significance after controlling for BMI. On the genus level (Fig. 2C), T2D patients had consistently increased proportions of *Libanicoccus*, *Lactobacillus, Collinsella*, *Senegalimassilia*, *Bifidobacterium* and *Slackia*, and reduced relative abundances of *Faecalibacterium* and *Oribacterium*. The relative abundance of *Collinsella* has been shown to correlate with elevated circulating insulin,^20^ increased gut permeability^21^ and altered bile acid metabolism^22^ that contribute to the pathophysiology of T2D. *Lactobacillus* represents one of the most discrepant signatures of the T2D gut microbiota, likely due to its high genomic diversity.^23^ Species were assigned to the ASVs that perfectly matched reference sequences, among which *Collinsella bouchesdurhonensis* and *Collinsella aerofaciens* were consistently enriched and *Faecalibacterium prausnitzii* depleted in T2D patients (Fig. 2D). The alternations in the bacterial taxa were expected to result in extensive changes in the metabolic potential of the gut microbiota, with 20 out of 27 KEGG pathways categorized as “metabolism” (Fig. 2F). Notably, the modules related to carbohydrate metabolism (i.e., inositol phosphate metabolism, galactose metabolism, glycolysis/gluconeogenesis, amino sugar and nucleotide sugar metabolism), amino acid-related metabolism (i.e., phosphonate and phosphinate metabolism, tyrosine metabolism, glutathione metabolism and selenocompound metabolism) and lipid metabolism (i.e., secondary bile acid biosynthesis and glycerolipid metabolism) were significantly more abundant in T2D patients (Fig. 2F). The bacterial phosphotransferase system (PTS), which mediates uptake of multiple sugars from the environment,^24^ was predicted to be higher in its capacity appreciably in the T2D gut microbiota (Fig. 2F). This, and perturbations in the functional modules related to amino acid metabolism, concord with the findings from the previous studies in Indian-Danish prediabetic and T2D patients.^11, 25^ We next investigated ecological measures of the gut microbiota in the control and case group. There was no difference in observed richness, but Shannon’s diversity and eubacterial density were drastically reduced in T2D patients, suggesting a strongly aberrant community (P < 0.001, Fig. 2G).

**Figure 2.**
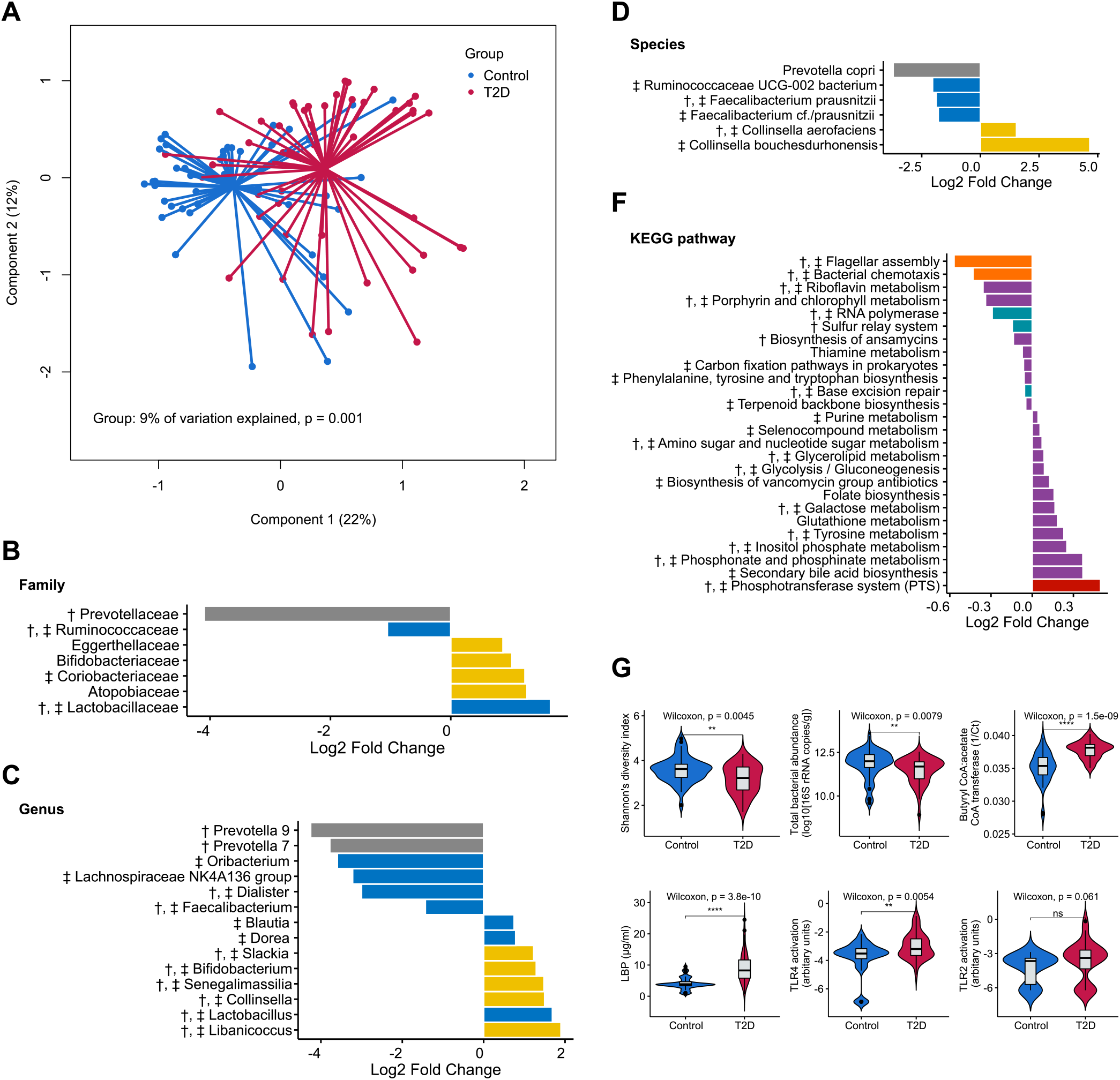
(**A**) Principal coordinates analysis (PCoA) plot based on the Bray-Curtis dissimilarity matrix showing the difference in the gut microbiota between Pakistanis without (blue) and with (red) T2D (P = 0.001, PERMANOVA). Differentially abundant bacterial **(B)** families **(C)** genera **(D)** species **(F)** KEGG pathways (level 3) are visualized by divergent bar plots. Only statistically significant results are shown (FDR-P < 0.05). Log2 fold change was calculated using controls as the reference group. Bacterial taxa are colored according to their respective phyla (Bacteroidetes in grey, Actinobacteria in blue, and Firmicutes in yellow). KEGG pathways are colored at level 1 (Environmental Information Processing in red, Metabolism in purple, Genetic Information Processing in teal, and Cellular Processes in orange). The differential bacterial taxa and KEGG pathways that remain significant after controlling for BMI or diet type are marked with ‡ and †, respectively. **(G)** Violin plots (a combination of the box plot with a kernel density plot) showing various measurements pertaining to gut microbiota ecology and microbial mediators. The center line denotes the median, the boxes cover the 25th and 75th percentiles, and the whiskers extend to the most extreme data point, which is no more than 1.5 times the length of the box away from the box. Points outside the whiskers represent outlier samples. Significance is calculated using the Wilcoxon rank-sum test. **** P < 0.0001; *** P < 0.001; ** P < 0.01; * P < 0.05; “ns” P > 0.05.

We additionally profiled the fungal communities using ITS1 sequencing, generating 7560 ± 785 quality-filtered ITS1 sequences per sample. Nevertheless, the majority of the pre-processed sequences failed to be taxonomically annotated, leaving only 28 participants with gut fungal profiles (control = 11, T2D = 17). The large fraction of unannotated reads suggests the existence of novel species from the Pakistanis not yet in sequence repositories. No difference in fungal β-diversity was found between the 28 participants with and without T2D (Fig. S1A). While the taxonomic composition was highly variable (Fig. S1B), *Candida sake, Teunomyces, and Candida akabanensis* were identified as significantly enriched in the patients with T2D (all FDR-P < 0.05, Fig. S1C).

We next assessed the molecular links between the gut microbiota and T2D, specifically butyrate as a major SCFA and PAMPs that have been suggested to be the cardinal microbial mediators in metabolic disorders.^1^ Butyrate production capacity estimated by quantifying the butyryl-CoA:acetate CoA-transferase gene (responsible for a major route for butyrate production in bacteria) was unexpectedly higher in T2D patients (P < 0.001, Fig. 2G). An increased influx of PAMPs in circulation, especially LPS (a component of the Gram-negative bacterial outer membrane), has been linked to inflammation and impaired glucose metabolism through activation of TLR4 or TLR2-dependent signaling.^26^ Thus, we measured the potential of plasma samples to activate innate immunity receptors TLR4 (receptor for LPS) and TLR2 (receptor for a variety of Gram-negative and Gram-positive bacterial products). The plasma samples from T2D patients induced significantly higher levels of TLR4 activation (P < 0.001, Fig. 2G), and elevated levels of TLR2 activation by trend (P = 0.061, Fig. 2G). For LPS-TLR4 signaling, LPS-binding protein (LBP) attaches LPS and presents it to CD14 to initiate TLR4 activation; we subsequently quantified the concentrations of plasma LBP and found increased levels of LBP in T2D patients (P < 0.001, Fig. 2G), mirroring the findings on TLR4 activation. Taken together, these findings suggest that Pakistani T2D patients possess an increased potential for butyrate production in the gut microbiota and engage in elevated TLR-signaling in circulation.

Since approximately half of the patients with T2D were prescribed with metformin in our cohort (Table 1), we conducted sub-group analysis for the patients to identify potential effects of metformin on the gut microbiota. Consequently, no significant differences in the abovementioned taxa and measurements were identified between the groups (Fig. S2), except lower levels of plasma LBP in those treated with metformin (P = 0.02, Fig. S2B).

### Correlations between metabolic parameters, bacterial taxonomic features and microbial mediators

We assessed all triplets of pairwise interactions between host metabolic phenotypes, bacterial genera and microbial mediators using Spearman correlations (requiring FDR < 0.05 in each comparison of two data spaces). Figure 3 shows a chord diagram constructed from these data, where the network largely reflects our observations mentioned previously (Fig. 2B-G). In general, the genera identified as enriched in T2D were positively associated with plasma LBP and HbA1c, whereas the genera belonging to the *Prevotellaceae* family were negatively associated with LBP, HbA1c and TG. Only two correlations were identified between TLR4 activation and bacterial taxa i.e., *Faecalibacterium* (reduced in T2D patients) that was negatively associated and *Libanicoccus* (enriched in T2D patients) positively associated with TLR4 activation (Fig. 5).

**Figure 3.**
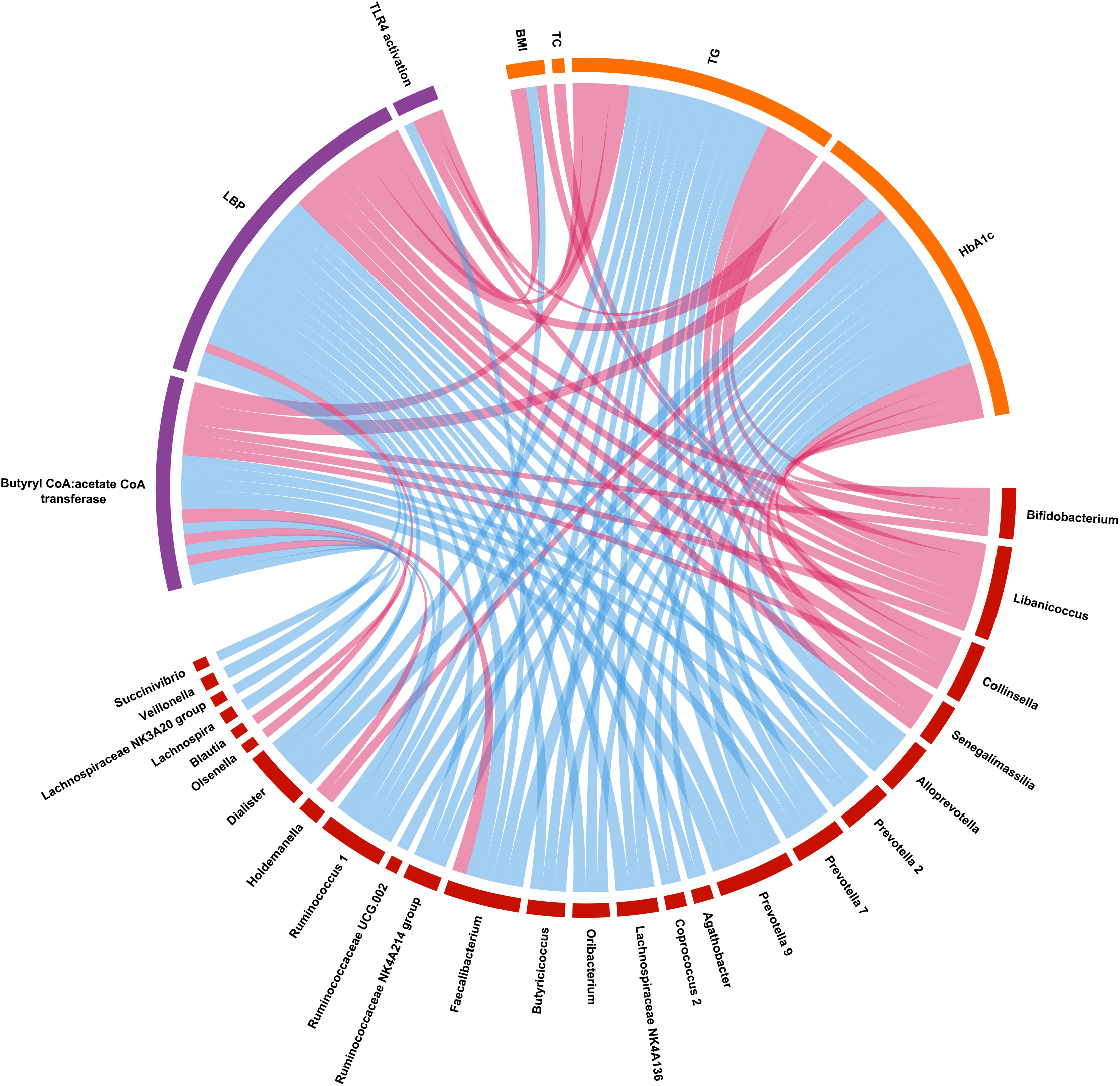
A Chord diagram visualizes the significant interrelation between host metabolic characteristics (cells in orange) and microbiota taxonomic features (cells in burgundy), and microbial mediators (cells in purple). Features are shown that form triplets of phenotype, microbial and microbial molecule-related variables where at least two of three correlations are significant (Spearman FDR-P < 0.05). Color of the connectors indicates positive (in red) or negative (in blue) Spearman’s rho values. BMI, body mass index; TC, total cholesterol; TG, triglycerides; HbA1c, hemoglobin A1c; LBP, lipopolysaccharide binding protein; TLR4, Toll-like receptor 4.

### Classification models of T2D based on bacterial taxonomic signatures and their cross-study portability

Varying and sometimes contrasting microbiota signatures of T2D have been documented in different cohorts and populations,^23, 27^ and their cross-study portability as biomarkers has been questioned.^11^ Given that we observed considerable taxonomic differences in the microbiota between the controls and T2D patients, many of which unreported previously e.g., *Libanicoccus* and *Slackia*, a random forest (RF) classifier model was constructed using the bacterial genera detected in our cohort (Table S3) that could specifically identify T2D patients with high overall accuracy (area under curve; AUC = 0.8684, Fig. 4A). This RF classifier was trained excluding the patients prescribed with metformin to avoid any unobserved effects of metformin. Next, we performed study-to-study model transfer (model trained for each study independently and their predictive performance evaluated on the other datasets) to systematically assess cross-study generalization of T2D microbiota signatures. The RF models in other cohorts were trained without excluding those treated with metformin, as the records of medication were unavailable on the individual level. Nevertheless, population-specific effects were evident, where the extrapolation of T2D classifiers performed satisfactorily among the Pakistanis, and to a lesser extent, the Sudanese (median AUC = 0.71, range = 0.62 – 1.0; Fig. 4B); relatively poor discriminatory power was pervasive for other populations (AUC < 0.7, Fig. 4B). To understand the population-specific discrimination accuracy, we applied *DESeq2* across different studies/cohorts to top 10 predictive microbial features extracted from the RF model trained on our cohort (Fig. 4C). The distribution of these features between controls and T2D patients was largely congruent among the two Pakistani and the Sudanese cohorts, but inconsistent or discordant at times in other cohorts (Fig. 4C). Of note, *Prevotella* 9, a highly discordant signature between the Pakistani and Chinese cohorts, was dominated by different clades of *Prevotella copri*; clade A was predominately present in the Chinese, while clades B and C were exclusively found in the Pakistanis (Table S2). Interestingly, *Libanicoccus* and *Slackia* appear to be the taxonomic signatures of T2D specific to Pakistanis, and the two taxa were prevalent in Pakistanis (Fig. S3). *Dialister* as a contrasting signature between the two Pakistani cohorts belongs to the few bistable taxa that differ in bimodal distribution patterns across cohorts, potentially indicating underlying alternative states associated with host factors.^29^ This cohort-specific bimodality of *Dialister* was observed between the two Pakistani cohorts (Fig. S4), suggesting the presence of cohort-specific characteristics (e.g., medical conditions unaccounted for in the two studies) in the same population that contributed to the contrasting signature. Of note, some difference between the two Pakistani cohorts of healthy controls was evident in the β-diversity analysis (Fig. 1A).

**Figure 4.**
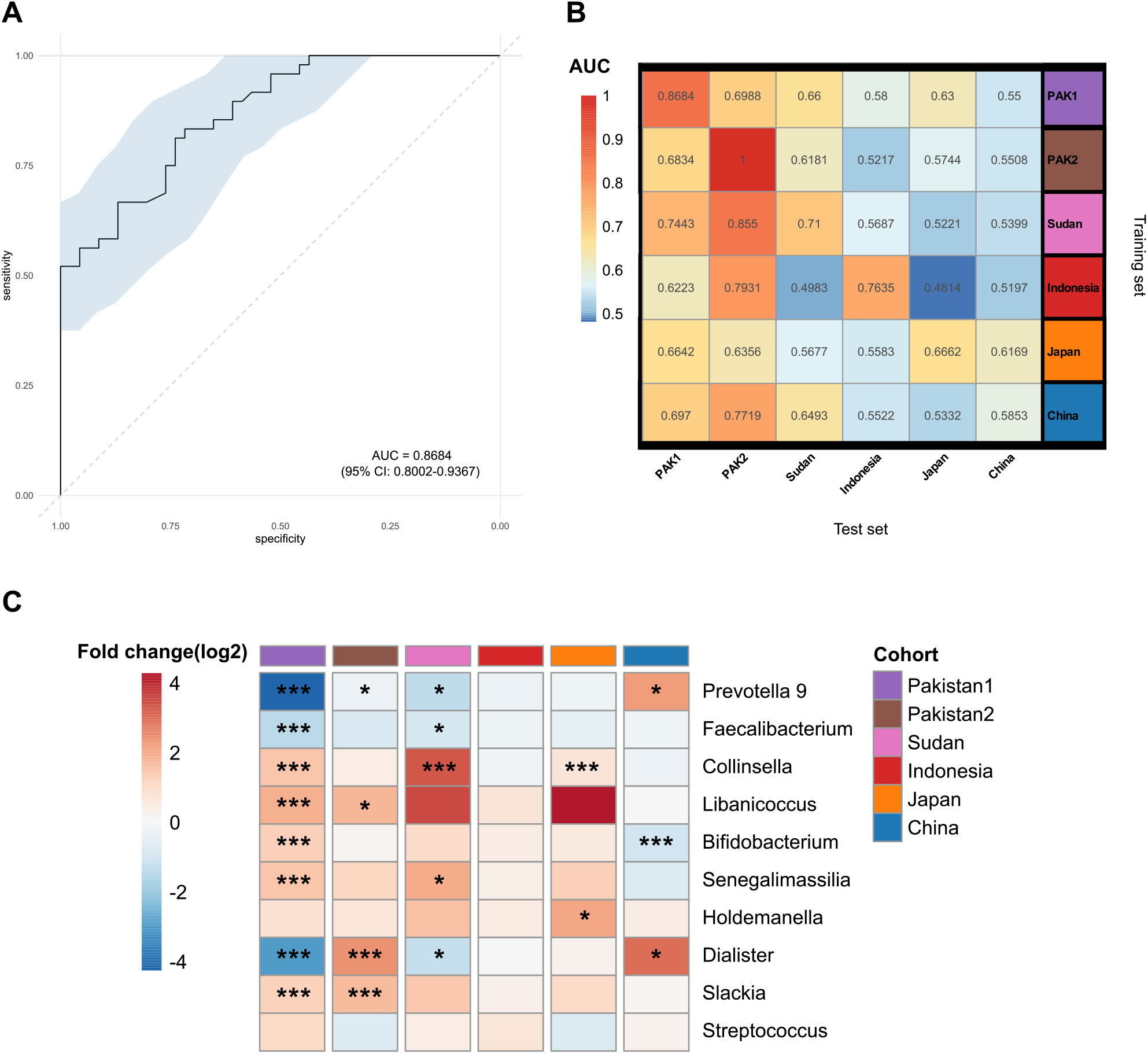
**(A)** A receiver operating characteristic (ROC) curve evaluating the ability to distinguish T2D cases from controls using a random forest classifier trained on bacterial genus-level abundances, achieving an area under ROC curve (AUC) value of 86.84% with 95%□CI of 80.02% to 93.67%. **(B)** T2D classification accuracy resulting from cross-validation within each study i.e., the test and training set are the same study (the boxed along the diagonal) and study-to-study model transfer (external validations off the diagonal) as measured by the AUC for the classification models trained on genus-level abundances. The color of the scale bar on the right represents the AUC value. **(C)** Abundance distribution of top 10 taxonomic features with the highest discriminatory power extracted from the random forest model trained on the present cohort (PAK1/Pakistan1). Log2 fold change was calculated using controls as the reference group and significance was determined by *DESeq2*. **** P < 0.0001; *** P < 0.001; ** P < 0.01; * P < 0.05.

## Discussion

The widespread changes in lifestyle and diet due to rapid urbanization in Pakistan over the last two decades have been linked to increased incidence of non-communicable diseases.^30, 31^ Urbanization and industrialization may remodel a population’s gut microbiota via multiple exposures, such as Westernization of diet, increased antibiotic use, pollution, improved or deteriorated hygiene status and early-life microbial exposure.^32, 33^ These selective forces could deviate the gut microbiota further from its ancestral state via intergenerational transmission.^34^ In some studies, the microbiota associated with humans in the industrialized world has been characterized by progressive disappearance of the *Prevotellaceae, Spirochaetaceae* and *Succinivibrionaceae* families, and increased abundance and prevalence of *Akkermansia* and *Bacteroides*.^2, 18^ Our results recapitulate these observations and place the urban Pakistani gut microbiota into the transitional or non-industrialized category. The persistence of non-industrialized microbiota signatures is in agreement with previous reports on transitioning populations or communities, such as first-generation migrants in the Netherlands,^8^ Irish Travellers enforced to abandon nomadism,^35^ and urban Nigerians.^36^ In contrast, a study about Thai migrants moving to the United States claimed that the microbiota rapidly assumes the structure of the new place of residence.^37^ While the degree and timeframe to which the microbiota signatures persist following lifestyle modernization remains an open question, transitioning from the non-industrialized to industrialized gut microbiota has been suggested to negatively affect metabolic health.^35, 37^ It remains unclear what caused the lack of detectable levels of *Akkermansia* in our Pakistani cohort and other non-industrialized populations. Nevertheless, it is possible that the absence of *Akkermansia* in the urban Pakistani adults is maladaptive, as this bacterium provides ecosystem services needed to maintain metabolic homeostasis especially in humans on the Western diet.^1^ Further research therefore is warranted to establish the causality between the transitioning gut microbiota and worsening metabolic health in developing countries.

Regarding the population-specific features of the Pakistani gut microbiota, our results are in agreement with a pilot study in a cohort of 32 urban Pakistani adults targeting the V4 region of the 16S rRNA gene.^14^ For example, the enrichment of *Bifidobacterium* and *Lactobacillus* is likely attributable to the frequent consumption of fermented foods in Pakistan;^14^ the occasionally high loads of *Enterobacteriaceae* and *Streptococcaceae* that contain several opportunistic pathogens likely result from deteriorated water and hygiene conditions during galloping urbanization. Data from our comparative analysis also reveals the high prevalence and abundance of *Prevotella* 9 in Pakistanis dominated by the non-Westernized clades of *Prevotella copri*, commonly thought to associate with plant-rich diets that are abundant in carbohydrates and fibers. The Pakistani diet is particularly low in intakes of fiber-rich fruits, vegetables, nuts and whole grains on the global scale.^38^ Also, there was no difference in the relative abundance of *Prevotella* between the Pakistani participants who consumed plant-based or meat-based diets (Fig. S5). Thus, the dominance of *Prevotella* in Pakistanis may have been promoted by non-dietary factors. This is supported by the study in Irish Travellers, a historically nomadic ethnic group of European ancestry, who consume a Western-like diet rich in fat and protein with little fiber intake.^35^ *P. copri* was found to be enriched to the same levels of other non-industrialized populations in a subgroup of Irish Travellers who maintained traditional lifestyles and living conditions, while no difference in diet was found between the subgroups of Irish Travellers.^35^ A recent study in rural Gambian infants found that *P. copri* rises to dominance rapidly in the first year of birth and remains the most abundant species during the remainder of early childhood,^39^ suggesting its establishment is strongly affected by early-life factors in *Prevotella*-rich populations. In addition to longer and more common exclusively breastfeeding, most weaning infants in Pakistan are initially given unprocessed foods as opposed to convenience baby foods given to infants in the Western world;^40^ solid foods introduced to most Pakistani infants are predominantly plant-based and family foods,^40^ particularly those made from roots and tubers that constitute a rich source of resistant starch.^41^ Taken together, future work should address the kinetics of colonization and development of *Prevotella* in relation to various dietary and non-dietary exposures in Pakistani infants.

As expected, both discrepancies and commonalities are noted regarding the taxonomic signatures of T2D between our findings and previous studies in various populations. The depletion of anti-inflammatory *Faecalibacterium prausnitzii* in T2D has been relatively consistent across studies.^23, 25^ Here we show a negative correlation between the abundance of *Faecalibacterium* in the fecal microbiota and TLR4 activation by the plasma, recently also reported for gut epithelium in an *in vitro* study.^42^ The enrichment of several members of the *Coriobacteriia* class (i.e., *Libanicoccus, Senegalimassilia, Slackia*) in Pakistani T2D patients, uncommon in the Western populations, appears to be novel population-specific signatures. *Libanicoccus massiliensis* was isolated from feces of a healthy Congolese pygmy woman in 2018,^43^ but has since been proposed to be reclassified as a member of the *Parolsenella* genus.^44^ *Slackia* has been sparsely described in the industrialized world, but relatively abundant in traditional societies e.g., rural Papua New Guineans.^45^ Little is known about *Libanicoccus*/*Parolsenella* as a newly isolated bacterium. A recent re-analysis of the studies in experimental autoimmune encephalomyelitis (EAE), the animal model of multiple sclerosis, associated *Parolsenella catena* with the development of pathology in marmosets;^46^ a metagenomic study in rural China reported that *Libanicoccus* was associated with arterial plaque buildup.^47^ The positive relationship between *Libanicoccus*/*Parolsenella* and TLR4 activation shown in the present study suggests a pro-inflammatory state in T2D potentially mediated by this bacterium.

Importantly, the abovementioned taxa belonging to *Coriobacteriia* are closely related, since the genome-based boundaries between its members are tenuous.^4, 44^ Almeida *et al*. reported that a large number of unclassified near-complete metagenomic species (i.e., potential new families and/or genera), most frequently assigned to the *Coriobacteriaceae* family, can be found in the gut microbiota data outside North America and Europe.^4^ Thus, our study highlights the importance of sampling underrepresented populations and regions to uncover novel microbes relevant for human health and improve the applicability of microbiota-based diagnostic strategies. The latter is further underscored by our findings on T2D classification models, where the discriminative power fell short when extrapolating a classification model to other populations. As the discriminative power of machine learning-based classification models is heavily influenced by the presence/absence of specific microbial taxa,^48^ additional variability from a greater pool of heterogeneous samples from various populations could enable classifiers to capture specific signatures while minimizing overfitting on idiosyncrasies of a single population. The potential of this strategy in improving microbiota-based disease classification has been demonstrated in recent proof-of-concept studies.^27, 49^

The patterns of *Prevotella* and *Bifidobacterium* in prediabetes and T2D are highly inconsistent across cohorts as shown in our and other studies.^25, 50, 51^ Higher abundance of *P. copri* in the gut microbiota has been suggested to be both conducive and detrimental to glucose homeostasis and host metabolism in different studies, and its role in human health is still under investigation.^52^ Tett *et al*. found no association between the prevalence and abundance of the four *P. copri* clades and the etiology of several diseases, including T2D.^19^ However, the two cohorts of T2D included in the Tett study are from the Chinese and European populations that generally have low prevalence of *P. copri* dominated by clade A. Therefore, the potential clade-specific effects of *P. copri* in T2D require further investigation in *Prevotella*-rich populations with non-Westernized *P. copri* clades using metagenomic sequencing. Additionally, we propose a scenario where some of the observed microbiota signatures in Pakistani T2D patients, namely the reduction in *Prevotella* and expansion in *Bifidobacterium* and butyrate production capacity, are reflective of their transitioning gut microbial ecosystem toward the industrialized one. Supporting this, the microbiota profiling of non-industrialized/non-urbanized African communities suggests a different ecological layout supporting SCFA production with little or no contribution from *Bifidobacterium* and typical butyrate producers,^36, 53^ corresponding to their “high-propionate and low-butyrate” SCFA profile.^53^ In general, the effects of *Bifidobacterium* and butyrate in promoting human health have been understudied in non-Western populations, where human-microbe associations may need to be interpreted in an entirely different context.^36^

A growing body of studies claims to have identified increased microbial components (e.g., LPS) in T2D, potentially contributing to its pathogenesis.^54^ These studies have nonetheless been challenged by various issues, including high technical variability and inability to differentiate stimulatory and inhibitory forms of LPS.^55, 56^ By directly quantifying the ability of plasma samples to elicit TLR-signaling, here we corroborated the elevated levels of circulating pro-inflammatory mediators in T2D patients. To our knowledge, this approach has not been applied to human blood samples. It is worth noting that other non-microbially derived molecules can also contribute to the inflammatory response by activating TLR4 signaling,^57^ which may partly explain few associations between the gut microbiota and TLR4 activation. Importantly, the gut microbial signatures do not necessarily translate to pro-inflammatory potential, which can be conceived as a net effect of pro-inflammatory microbial mediators, translocation rate of the mediators and host’s ability to handle them.

Our comparative analysis is limited by sample size owing to missing metadata or restricted data availability in many published studies;^58^ the available metadata for all the 944 samples used in the present study has been included in Table S4. We were nonetheless able to recapitulate critical observations made by previous studies e.g., the traits of the non-industrialized gut microbiota.^18^ Moreover, the unique characteristics of the Pakistani gut microbiota were further validated in an independent cohort. On the other hand, our study focusing on urbanized Pakistanis of a similar social background was unable to fully capture the full diversity of Pakistan. To substantiate our findings and hypothesis on the transitioning gut microbiota, future large studies should expand to peri-urban and rural communities. Although we did not observe the effects of metformin on the gut microbiota previously reported in Chinese, Japanese and European cohorts of T2D patients,^59^ the beneficial effects of metformin have been suggested to be partially mediated by the gut microbes, such as *P. copri*^60^ and *Akkermansia muciniphila*^61^ that have different abundance and prevalence distributions in Pakistanis. Therefore, further research is warranted to assess the potential contribution of the gut microbiota to the variability in the therapeutic response of metformin in the Pakistani population.

In conclusion, we show that the urbanized Pakistanis’ gut microbiota retains non-industrialized features with several distinctive traits. These unique microbiota characteristics extend to T2D-associated signatures, potentially reflecting transitioning gut microbial ecosystems and contributing to pro-inflammatory states, which hold significance for diagnostic and therapeutic applications.

## Methods

### Study participants and sample collection

The present case-control study was performed during the last quarter of 2021 at the National Institute of Health, Islamabad, Pakistan. Adults (> 18 years old) with confirmed type 2 diabetes according to the American Diabetes Association (ADA) criteria^62^ (n = 48) and age- and sex-matched healthy controls (n = 46) were recruited via primary and occupational health care providers to the present study. All the participants are currently residing in Islamabad and Rawalpindi that originally belong to rural regions of Punjab and Khyber Pakhtunkhwa provinces. Exclusion criteria included type 1 diabetes, significant diseases including cardiovascular, liver, or kidney disease, malignancy, bariatric or any major surgical procedure in the previous 3 months, gastrointestinal diseases or symptoms of constipation or diarrhea, pregnancy or breastfeeding, use of antibiotics in the previous 3 months, being underweight (BMI < 18.5 kg/m^2^), and alcohol consumption.

All participants underwent anthropometric measurements at the clinic, and medical/drug history was documented for individuals with T2D. All participants also completed simplified food frequency questionnaires. The questionnaires were further converted to four types of diet for microbiota analysis (Table S5); this reductionist approach was mainly for convenience, but also recent studies suggest that dietary patterns and food groups associate more strongly with microbiota composition than conventional nutrients.^63^ Participants were informed of the need for fresh samples, and thus stool samples were collected in sterile containers provided to the participants and immediately transferred to −80°C freezer within 1 hour of defecation. Blood samples were analyzed for HbA1c and for lipid profiles (total cholesterol, triglycerides, LDL and HDL) by an automated analyzer (Cobas Integra 700; Hoffman-La Roche, Basel, Switzerland). The rest of the plasma samples were stored at −20°C until further analysis.

The healthy Finnish controls were previously recruited as part of a dietary intervention trial investigating the effects of partly replacing animal proteins with plant proteins on health; the inclusion and exclusion criteria have been described elsewhere.^64^ The fecal samples collected at baseline from 50 Finnish adults that were age-, sex- and BMI-matched with the Pakistani controls were included in the present study.

The study was carried out in accordance with the Declaration of Helsinki. The protocols were approved by the Ethics Review Committee (ERC) of the National Institute of Health, Islamabad, and the Medical Ethical Committees of the Hospital District of Helsinki and Uusimaa and HUCH. All subjects gave written informed consents.

### Publicly available datasets from other cohorts

To compare the Pakistan gut microbiota with that of other cohorts or populations, we systemically searched for publicly available 16S rRNA gene amplicon (targeting the V3-V4 region) datasets of T2D case-control studies with available metadata or identification of T2D status. Sequences from five published studies were subsequently downloaded from the National Center for Biotechnology Information and the DNA Data Bank of Japan: PRJNA661673 (China),^65^ PRJNA766337 (Japan),^66^ PRJDB9293 (Indonesia),^67^ PRJNA588353 (Sudan),^68^ and PRJNA554535 (Pakistan; an independent cohort consisting of participants residing in Islamabad).^15^ An endogamous agriculturist Indian cohort of healthy adults (PRJNA399246) was also included,^69^ representative of a non-industrialized population from the same geographic region. The characteristics of the included cohorts are summarized in Table S1.

### DNA extraction and sequencing of 16S rRNA gene and internal transcribed spacer-1 (ITS-1) amplicons

DNA was extracted from the Pakistani and Finnish participants’ fecal samples using the same procedure, Repeated Bead Beating (RBB) method,^70^ with the following modifications for automated DNA purification: Approximately 0.25 grams of fecal samples and 340 μL and 145 μL of lysis buffer was used on the first and second round of bead beating, respectively. Then, 200 μL of the clarified supernatant collected from the two bead beating rounds was used for DNA extraction with the Ambion Magmax™ −96 DNA Multi-Sample Kit (4413022, Thermo Fisher Scientific, USA) using the KingFisherTM Flex automated purification system (ThermoFisher Scientific, USA). DNA was quantified using Quanti-iT™ Pico Green dsDNA Assay (Invitrogen, San Diego, CA, USA). Library preparation and Illumina MiSeq sequencing of the hypervariable V3-V4 regions of the 16S rRNA gene using primers 341F/785R were performed as previously described.^71^ To characterize the gut fungal community, we performed ITS-1 sequencing for the Pakistani samples using a two-step PCR protocol described in detail elsewhere.^72^ PCR-amplicons of the ITS-1 region were generated using ITS1F and ITS2 primers.^72, 73^

### Quantification of butyrate production capacity and eubacterial quantitative PCR (qPCR)

The butyryl-CoA:acetate CoA-transferase gene and total bacterial abundance were quantified using fecal DNA with the degenerate primers BCoATscrF/R and the universal primers 331F/797R by qPCR, respectively. The qPCR assays have been described in detail previously^71, 74^ and were performed in triplicate on a BioRad C1000 Touch thermal cycler (BioRad, Hercules, CA) with HOT FIREPol^®^ EvaGreen^®^ qPCR Mix Plus (Solis BioDyne, Tartu, Estonia). For quantification of the butyryl-CoA:acetate CoA-transferase gene, the mean threshold cycle (Ct) per sample (after excluding triplicates with Ct values that differed > 0.5) was used as a proxy for the abundance of the target gene. For quantification of total eubacteria, the 10-log-fold standard curves ranging from 10^2^ to 10^7^ copies were produced using full-length amplicons of 16S rRNA gene of *Bifidobacterium longum* to convert the threshold cycle (Ct) values into the average estimates of target bacterial genomes present in 1 g of feces (copy numbers/g of wet feces) in the assays.^71^

### Data processing and statistical analysis

For the bacterial microbiota, demultiplexed reads after adaptor removal were processed using DADA2^75^ to generate amplicon sequence variants (ASVs). Taxonomic classification was performed using a naive Bayes classifier against the SILVA 132 reference database^76^ for all the included cohorts. Species assignment was performed using DADA2 by exact string matching against the SILVA species assignment training database.^75^ The 16S rRNA gene sequences for the Pakistani and Finnish samples from this study have been deposited in the European Nucleotide Archive (https://www.ebi.ac.uk/ena) under accession numbers PRJEB53017 and PRJEB53018, respectively. For the fungal microbiota, the ITS-1 sequences were pre-processed following the DADA2 ITS Pipeline Workflow (1.8) available on the official DADA2 homepage, and taxonomically annotated using UNIITE ver. 7.2 Dynamic Classifier as the reference database with the similarity threshold at 0.97.^77^ The ITS-1 sequences have been deposited in the European Nucleotide Archive (https://www.ebi.ac.uk/ena) under accession number PRJEB53019.

To infer the functional contribution of bacterial communities from 16S rRNA gene sequencing data, metagenome prediction was carried out using PICRUSt2 (Phylogenetic Investigation of Communities by Reconstruction of Unobserved States)^78^ evaluating KEGG (Kyoto Encyclopedia of Genes and Genomes) pathways.^79^

Differential abundance for bacterial and fungal taxa or KEGG pathways between case and control participants was identified with the *DESeq2* package.^80^ *DESeq2* employs a generalized linear model of counts based on a negative binomial distribution, scaled by a normalization factor that accounts for differences in sequencing depth between samples. Significance testing was then assessed using the Wald test. Non-count variables (anthropometric, biochemical and other measurements) were analyzed with the Wilcoxon signed-rank test, t-test or chi-square test depending on the data distribution.

Microbiota α-diversity (observed richness and Shannon diversity index) was estimated using the *vegan* package.^81^ Overall microbiota structure was assessed by principal coordinate analysis (PCoA) based on β-diversity computed using the Bray-Curtis dissimilarity matrix, representing the compositional dissimilarity between samples or groups. Significant differences between groups were tested using nonparametric multivariate analysis of variance (PERMANOVA).^81^ The associations between continuous or categorical variables and β-diversity were calculated using the *envfit* function in the *vegan* package,^81^ and P values were determined using 999 permutations. Associations between relative abundances of bacterial taxa and other measurements were assessed using Spearman’s correlation and visualized by the *R* packages *circilize*.^82^

For microbiota-based T2D classification, a random forest model (number of trees = 1000; *R* package *randomForest*) was built based on a list of bacterial genera detected in our Pakistani cohort (Table S3). The model performance was evaluated using repeated k-fold cross-validation (5-fold, 10 repetitions) implemented in the *caret* package^83^ and scored the predictive power in a receiver operating characteristic (ROC) analysis. The top 10 important input genera were subsequently ranked according to mean decrease in Gini. To assess how well the classifier trained on one cohort can be generalized to the other cohorts, study-to-study model transfer was performed as described previously^27^. Briefly, one random forest model was built based on the abovementioned list of input genera for each study/cohort and then applied to the other studies using the same parameters to generate area under curves (AUCs) for cross-applications.

Statistical analyses were performed with the statistical program R version 3.5.0 and RStudio version 0.99.903. P values were corrected for multiple comparisons by using the Benjamini-Hochberg procedure (FDR). P values and FDR-adjusted P-values < 0.05 were considered significant.

### Inference of clade of Prevotella copri ASVs

The reconstructed genome sequences of *Prevotella copri* by Tett *et al*.^19^ were collected and converted into a BLAST database using the BLAST+ package.^84^ The sequence of each ASV annotated as *Prevotella copri* was queried to the database using the *blastn* command, and the reconstructed genomes containing the ASV sequence (100% identify) were retrieved. The clades of these genomes were assigned as the clade of the ASV.

### Measurement of Toll-like receptor (TLR) activation and lipopolysaccharide binding protein (LBP)

To quantify potential microbial products in circulation, TLR activation capacity and levels of LBP in plasma samples were used as a proxy. The protocol using HEK-Blue™-hTLR2 and HEK-Blue™-hTLR4 reporter cell lines expressing human TLR2 and TLR4 (InvivoGen, San Diego, CA) was employed according to the manufacturer’s instructions. The HEK-Blue™ hTLR cells were grown for two passages with medium supplemented without selective antibiotics provided by the manufacturer, and then passaged in medium with selective antibiotics that was also used for the experiment. The assay was performed when cells were in passage 10-15 by adding approximately 10^5^ HEK-Blue™-hTLR2 cells and 1.4× 10^5^ HEK-Blue™-hTLR4 cells in 96-well plates containing 20 μl of plasma samples and incubated for 24 hours at 37 °C under an atmosphere of 5% CO_2_/95% air. Twenty microliters of the cell culture supernatants were added to 180 μl of the QUANTI-Blue substrate in a 96-well plate. The mixtures were then incubated at 37°C for 3 hours and secreted embryonic alkaline phosphatase levels were determined using a spectrophotometer at 630 nm. Lipoteichoic acid (Invivogen, LTA-BS; 1 μg/ml, 500 ng/ml, and 100 ng/ml), peptidoglycan (Invivogen, PGN-BS; 0.1 μg/ml, 5 μg/ml, 10 μg/ml,), and lipopolysaccharide (Sigma Aldrich, LPS, 1000 ng/ml, 100 ng/ml, 10 ng/ml) were used as positive controls, and cell culture medium served as a negative control. Due to reagent availability, the TLR2 experiment was performed for the samples derived from 50% participants whose age, sex, and BMI matched the entire cohort (control = 23, T2D = 24). Plasma LBP was measured by quantitative ELISA using human LBP DuoSet kits (R&D Systems, Minneapolis, MN) according to the manufacturer’s instructions.

BMI: body mass index
TC: total cholesterol
TG: triglycerides
HbA1c: hemoglobin A1c
T2D: type 2 diabetes mellitus
Met−: T2D patients prescribed with non-metformin hypoglycemic treatment
Met+: T2D patients prescribed with metformin.

## Data availability

The datasets generated in this study are available in the European Nucleotide Archive (ENA) repository, under accession no. PRJEB53017, PRJEB53018 and PRJEB53019. The associated metadata is available in Table S4. Supplemental data for this article can be accessed on publisher’s website.

## Authors’ contributions

Afshan Saleem: Conceptualization, Methodology, Formal analysis, Investigation, Resources,

Data Curation, Project administration, Writing - Original Draft, Visualization, Funding acquisition

Aamer Ikram: Conceptualization, Resources, Project administration, Writing - Review & Editing

Evgenia Dikareva: Methodology, Investigation, Writing - Review & Editing

Emilia Lahtinen: Methodology, Investigation, Writing - Review & Editing

Dollwin Matharu: Methodology, Investigation, Writing - Review & Editing, Supervision

Anne-Maria Pajari: Resources, Data Curation, Writing - Review & Editing

Willem M. de Vos: Methodology, Resources, Funding acquisition, Writing - Review & Editing

Fariha Hasan: Conceptualization, Resources, Project administration, Writing - Review & Editing

Anne Salonen: Conceptualization, Methodology, Resources, Writing - Review & Editing, Supervision, Funding acquisition

Ching Jian: Conceptualization, Methodology, Formal analysis, Investigation, Data Curation, Writing - Original Draft, Visualization, Supervision

