## Supplementary_Figures for "Unique Pakistani gut microbiota highlights population-specific microbiota signatures of type 2 diabetes mellitus"

### Figure S1

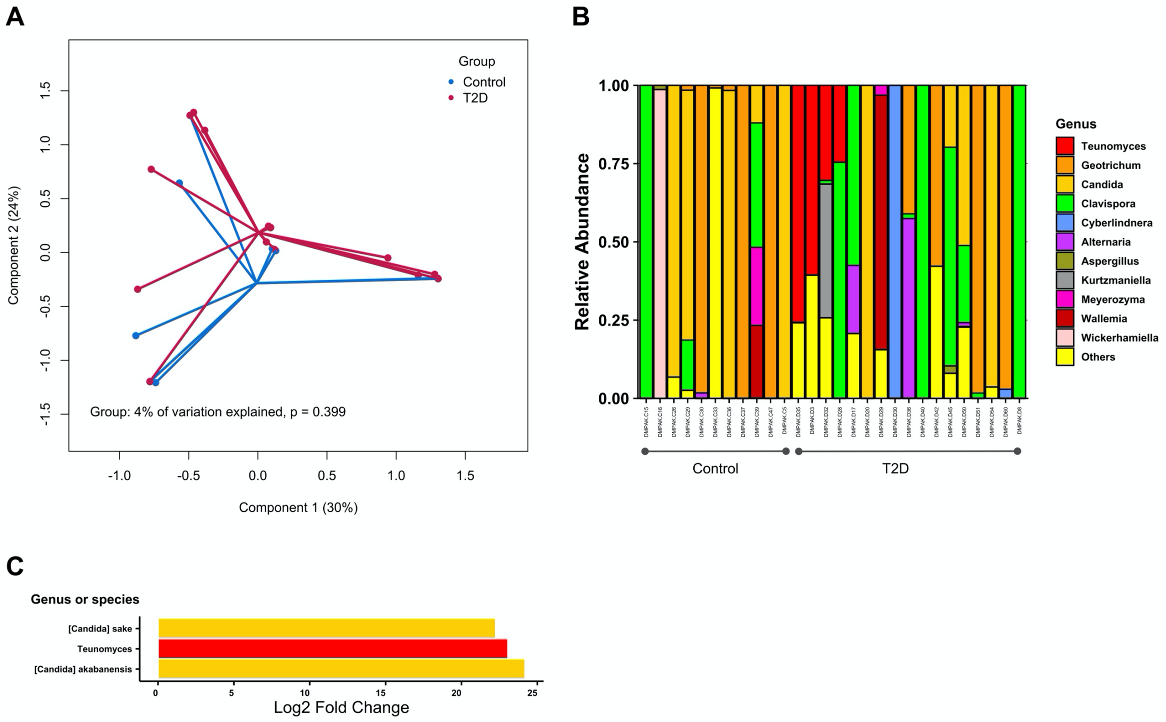


**Figure S1.** (**A**) Principal coordinate analysis (PCoA) of microbiota variation based on the Bray-Curtis dissimilarity matrix, showing no difference in the overall fungal microbiota between controls (blue) and T2D patients (red) (P = 0.399, PERMANOVA). (**B**) Stacked bar plots showing the composition of fungal genera. (**C**) Differentially abundant fungal genera and species between controls and T2D patients. Only statistically significant results are shown (FDR-P < 0.05). Log2 fold change was calculated using controls as the reference group.

**Figure S2**

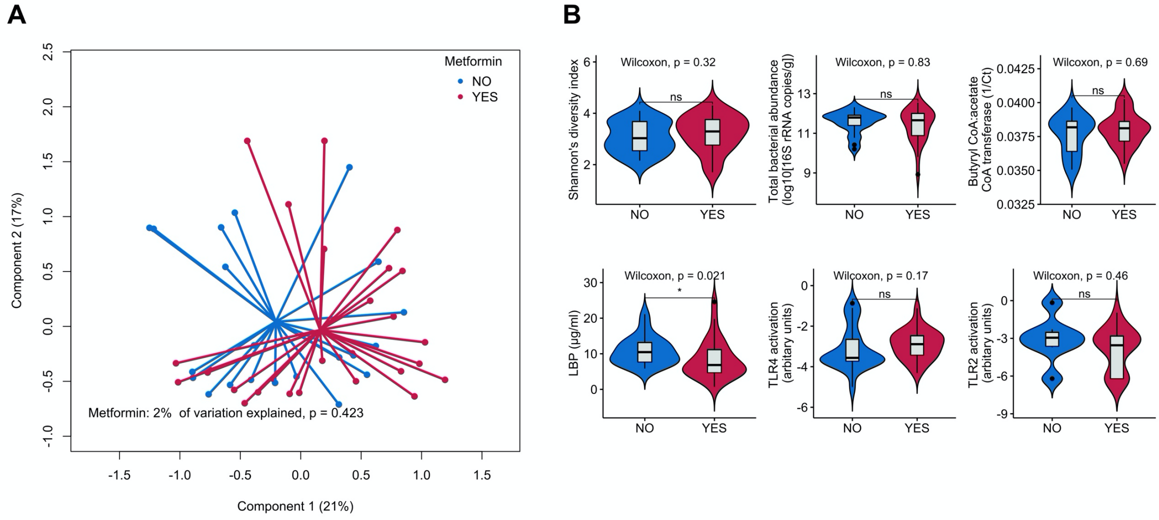


C

| Genus | baseMean | log2FoldChange | lfcSE | stat | pvalue | padj |
|---|---|---|---|---|---|---|
| Family XIII AD3011 group | 9.3566 | 2.0540 | 0.7556 | 2.7185 | 0.0066 | 0.4813 |
| *Bifidobacterium* | 1100.3311 | -0.9930 | 0.3726 | -2.6649 | 0.0077 | 0.4813 |
| *Ruminiclostridium* 6 | 59.1589 | -2.7877 | 1.0493 | -2.6567 | 0.0079 | 0.4813 |
| *Prevotella* 9 | 9.2291 | 2.2702 | 0.9571 | 2.3721 | 0.0177 | 0.8093 |
| *Fusobacterium* | 43.1613 | 3.1630 | 1.4137 | 2.2373 | 0.0253 | 0.8409 |
| *Negativibacillus* | 8.1796 | 2.6013 | 1.1806 | 2.2034 | 0.0276 | 0.8409 |

**Figure S2.** **(A)** Principal coordinate analysis (PCoA) of microbiota variation based on the Bray-Curtis dissimilarity matrix, showing no statistical difference in the overall gut microbiota between T2D patients treated without (blue) and with (red) metformin (P = 0.399, PERMANOVA **(B)** Comparison of ecological measures of the gut microbiota, butyrate production capacity of the gut microbiota, circulating LBP and TLR activation between patients with T2D prescribed with metformin or non-metformin hypoglycemic treatment. **(C)** Output of *DESeq*2 showing differentially abundant genera between T2D patients treated without and with metformin that had nominal P values < 0.05. The significant was lost after adjusting for multiple testing (all FDR-P > 0.05). Log2 fold change was calculated using non-metformin users as the reference group.

**Figure S3**

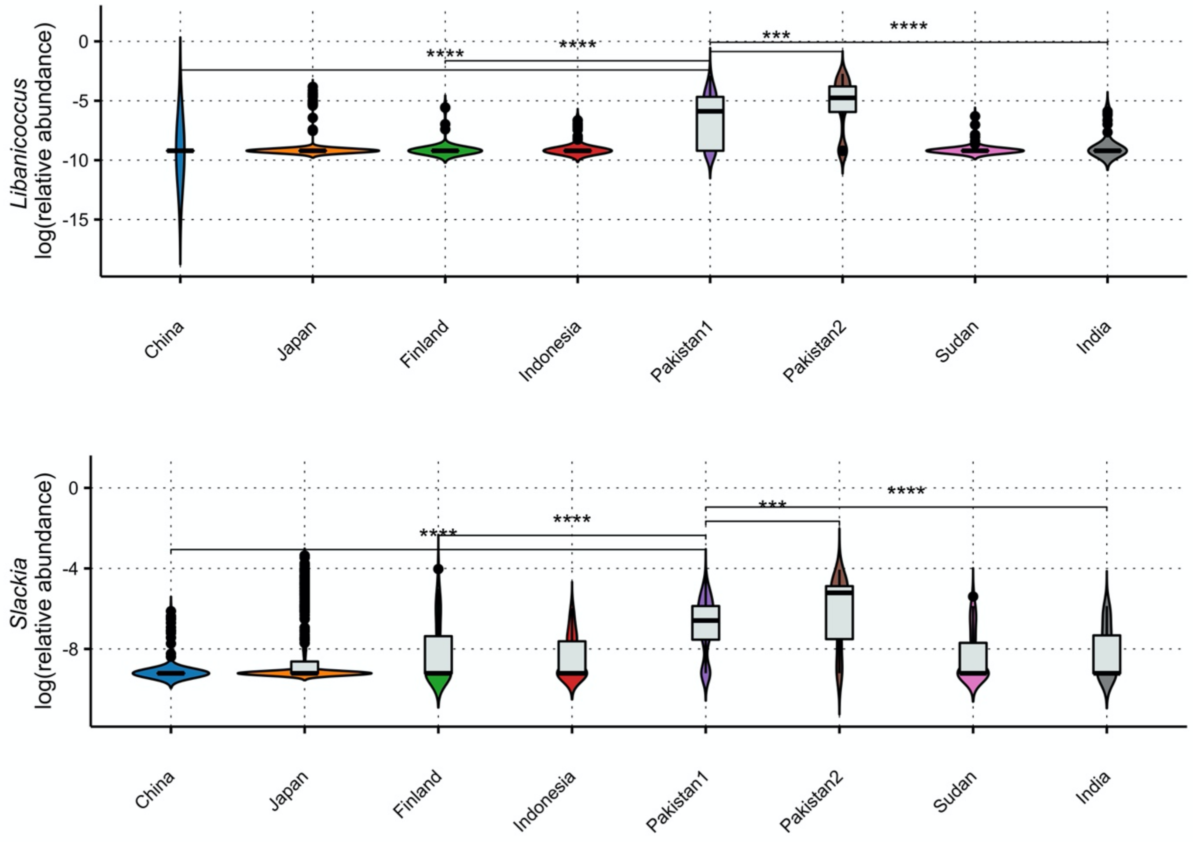


**Figure S3.** Violin plots (a combination of the box plot with a kernel density plot) showing the relative abundance of *Libanicoccus* and *Slackia* per each cohort. The center line denotes the median, the boxes cover the 25th and 75th percentiles, and the whiskers extend to the most extreme data point, which is no more than 1.5 times the length of the box away from the box. Points outside the whiskers represent outlier samples. Significance is calculated using the Wilcoxon rank-sum test. **** $P < 0.0001$; *** $P < 0.001$; ** $P < 0.01$; * $P < 0.05$.

**Figure S4**

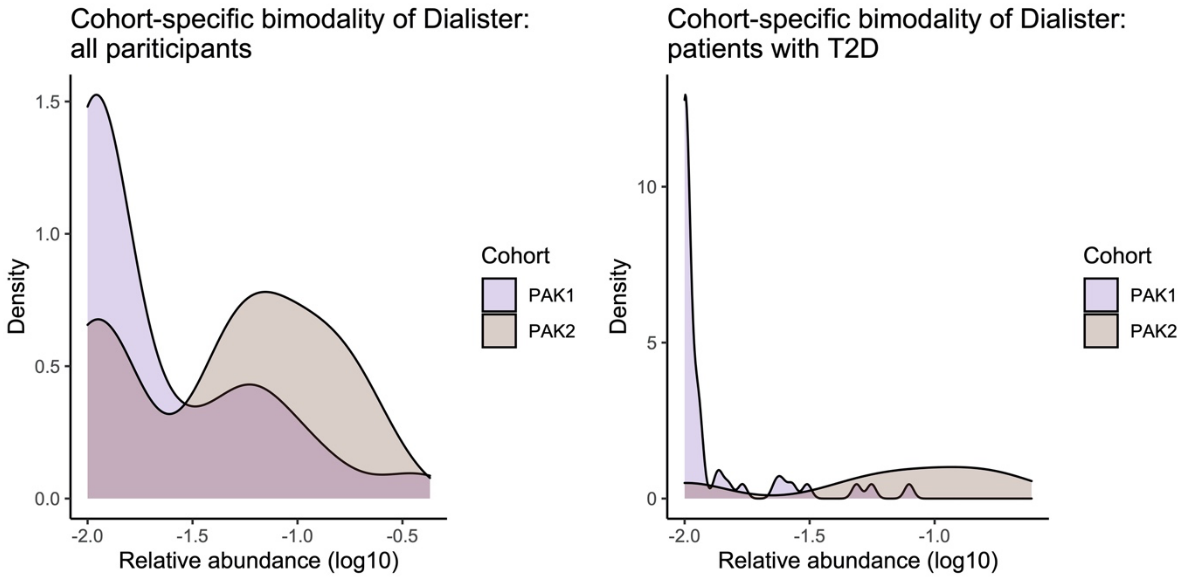


**Figure S4.** Observation density of *Dialister* in controls (left) and T2D patients (right).

**Figure S5**

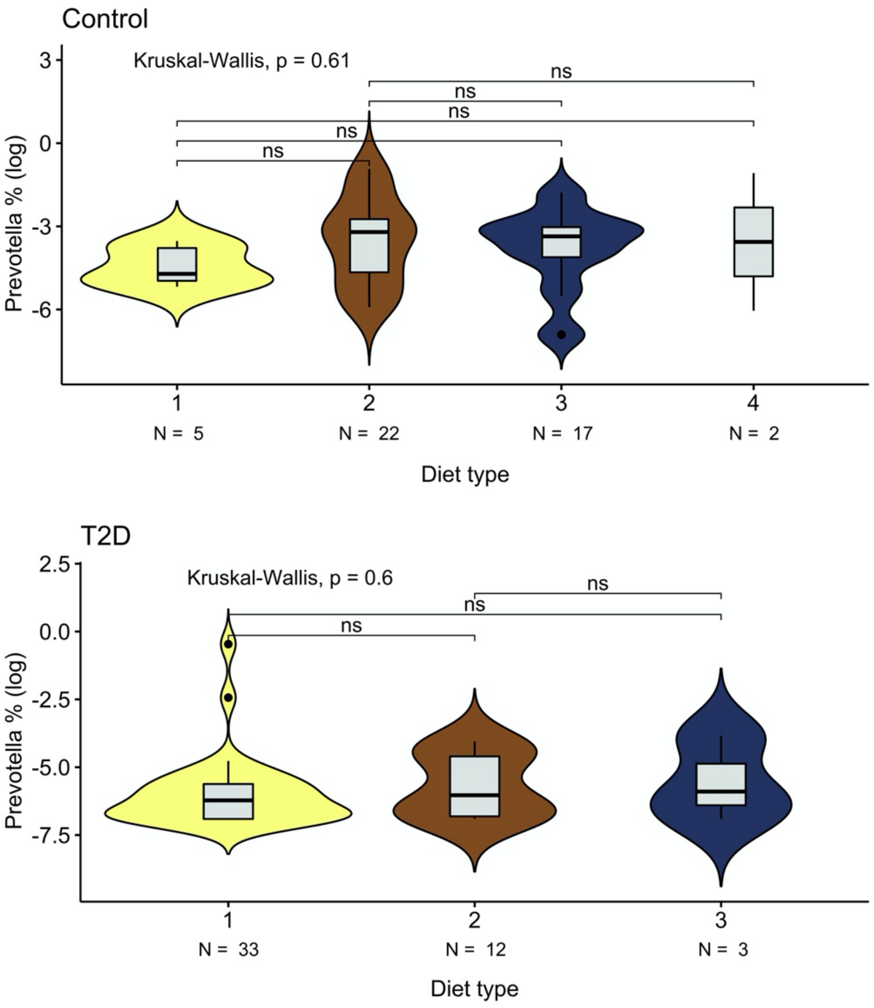


**Figure S5.** Violin plots (a combination of the box plot with a kernel density plot) showing relative abundance of *Prevotella* according to diet (Table S5). The center line denotes the median, the boxes cover the 25th and 75th percentiles, and the whiskers extend to the most extreme data point, which is no more than 1.5 times the length of the box away from the box. Points outside the whiskers represent outlier samples. Significance is calculated using the Wilcoxon rank-sum test. "ns", $P > 0.05$.
